# Characterization and Genomic Analysis of vB_CoeS_P1, the First *Vibrio hepatarius* Phage, and Definition of the New Family *Estovirdae*

**DOI:** 10.1101/2025.04.29.651204

**Authors:** P K Shruthi, Parvathi Ammini, Pradeep Ram Angia Sriram

## Abstract

A novel bacteriophage vB_CoeS_P1, isolated in the surface waters of the Cochin estuary of Kerala, India, was identified as the first phage known to infect *Vibrio hepatarius*. Transmission electron microscopy analysis revealed that the phage had a myovirus morphotype with an icosahedral head and a long contractile tail. Characterization experiments show that the phage was stable across a temperature range of 4°C to 60°C, a pH of 4 to 12, and a salinity range of 0.5 % to 16%. One-step growth curve analysis revealed a burst size of 30 PFU/Cell and a latent period of 20 minutes. The phage contains a double-stranded DNA with 136,927 bp and 34.93% GC content. The NCBI Blastn analysis reveals that vB_CoeS_P1 exhibits very low sequence similarity (1% query coverage and 77.82% identity) with Vb_VpaM_R16F (OP793884.1). It has 265 predicted ORFs, among them 57 have putative functions with no tRNA, antibiotic encoding genes, and resistance encoding genes. Three putative auxiliary metabolic genes were identified, encoding pyrophosphohydrolase enzyme (ORF 75), PhoH-like phosphate starvation-inducible (ORF 34), and Endonuclease V N-glycosylase UV repair enzyme (ORF 248). Phylogenetic analysis of conversed genes revealed that vB_CoeS_P1 is clustered with Vb_VpaM_R16F but diverged into a distinct clade. Further whole genome proteomic tree analysis, comparative genome analysis using average nucleotide identity (ANI), and network analysis confirmed that vB_CoeS_P1 contains a unique genetic and evolutionary architecture. These findings support the proposal of a new bacteriophage family *Estovirdae,* within the *Caudoviricetes* class.

## INTRODUCTION

Marine viruses represent a vast and large unexplored genetic reservoir on Earth. Most viruses in the marine ecosystem are phages, viruses that attack bacteria. Viruses exert substantial control over the abundance and diversity of their host organisms, accounting for the mortality of 25 to 50% of marine microbial biomass daily. Seawater is estimated to contain virus-like particles (VLPs) at an average concentration of 10^7^ VLPs per mL^-1^ (Wommack & Colwell, 2000). Phage-host interactions have a critical role in the oceanic biogeochemical balance. These interactions have both short-term and long-term effects. In the short term, the viral shunt redirects the flow of carbon by lysing microbial biomass, disrupting the transport of carbon to higher trophic levels (1). In the long term, the viral shuttle increases the flow of carbon to the sea floor via the biological pump, facilitating carbon sequestration (2).

Viral-mediated horizontal gene transfer (HGT) is a primary factor for functional and genetic diversity in microbial communities. It affects the host’s nutrition, defense, function, and adaptability (3). Marine phages also contain phage-encoded, host-derived metabolic genes, known as Auxiliary Metabolic Genes (AMGs), which help to enhance viral replication by upregulating the metabolic process in the host. Most viral AMGs encode proteins related with photosynthesis, carbon metabolism, translational regulation, NADH dehydrogenase function etc. So, phages can influence host diversity and metabolism directly or indirectly (Hurwitz & U’Ren, 2016).

In marine ecosystems, host-phage interactions are complex and shaped by various ecological and evolutionary hypotheses. One of the most prominent hypotheses is "killing the winner," where phage populations primarily infect dominant microbial spp. in the ecosystem, enabling less abundant microbial species to flourish and thus enhancing diversity within the microbial communities (5). In another hypothesis, called the “Piggyback-the-Winner" hypothesis, the phages adopt a lysogenic lifecycle and help the co-existence of the host and their phage without exerting immediate fatal pressure, allowing both to survive (6). The "King of the Mountain"(KoM) hypothesis emphasizes that the dominant host population is sustained in the ecosystem by developing resistance (through CRISPR and horizontal gene transfer) against phage-mediated lysis, and a balance between lysis and genetic diversity stabilizes the co-existence of both the phage and the host (7). This is supported by studies on SAR11 bacteria and their phages, which demonstrated that the high recombination enhances the chance of co-existence of both hosts and phages (8). Host-page coevolution is characterized by a successive cycle of adaptation and counter-adaptation, where host spp. evolves a resistant mechanism, and the phage develops mutations to overcome these defenses.

Vibrios are one of the most prevalent and ecologically significant bacteria in marine, estuarine, and freshwater ecosystems. These gram-negative, rod-shaped Proteobacteria fall under the class Gammaproteobacteria. *Vibrio hepatarius*, an important species in the genus Vibrios, was first isolated from white shrimp *Litopenaeus vannamei* in Ecuador. This facultatively anaerobic bacterium ferments glucose, sucrose, mannitol, and amygdalin. *Vibrio hepatarius* thrives in marine environments, particularly in estuarine and coastal waters (9). It also contributes to microbial communities within marine invertebrates, including shrimps, corals, and other crustaceans. *Vibrio hepatarius* has been identified in the coral *Porites lutea*, affected by white patch syndrome across geographically distant sampling locations in the western Indian Ocean reefs (10).

Within the scope of this study, we isolated and characterized a novel *Vibrio hepatarius* phage, vB_coes_P1, from the coastal waters of the Cochin estuary, and this is the first phage of *Vibrio hepatarius* to be reported to our knowledge. The International Committee on Taxonomy of Viruses (ICTV) eliminated the morphology-based families of phages, Myoviridae, Podoviridae, and Siphoviridae In 2021, the order Caudovirales was removed and replaced with the class *Caudoviricetes* (Turner et al., 2023). Advancements in genome sequencing and bioinformatic tools provide a refined and flexible approach to virus classification. The new methods classify viruses based on their phylogenetic and evolutionary relationships rather than depending on their phenotypic traits (Turner et al., 2021). In this context, each newly discovered novel phage contributes knowledge to the expanding viral diversity databases and provides insight into the unexplored fields of host-phage interaction studies. Genomic and phylogenetic analyses support the proposal of a new viral family for vB_coes_P1. This study not only enhances scientific insight into phage biology and genomic architecture but also serves as an excellent model for exploring marine host-phage interactions, and its unique evolutionary lineage sheds light on the origins and evolution of viral diversity.

## Materials and Methods

### Bacterial stain isolation and purification

The water sample was collected from the Cochin estuary (9°59’50.9"N 76°16’13.9"E) on November 13, 2023. Serial dilutions were made up to 10^-6^ in physiological saline and from each dilution, a 100 μL sample was spread on Zobell Marine Agar (ZMA) plates and incubated at 27°C. After incubation, sixteen distinct bacterial colonies with different morphological characteristics were obtained. These bacterial colonies were purified four to five times until a pure culture was obtained.

### Identification of host bacteria

Among the sixteen isolates, only one isolate produced plaque with enriched water samples. This specific isolate was chosen for further purification and identification. The Phenol-chloroform-isopropanol protocol was used for the isolation of bacterial DNA. Polymerase Chain Reaction (PCR) was used to amplify the 16S ribosomal RNA gene, followed by sequencing. The sequence was queried against NCBI BLASTn to identify the strain. To classify the bacterial strain taxonomically, a phylogenetic tree was generated using the neighbor-joining approach in MEGA 11 software, with 1000 bootstrap replicates.

### Isolation of phage from water

Phage isolation and purification followed the standard virus enrichment method and a double-agar overlay technique (11). Initially, the water sample was serially filtered through a mesh (20 nm) and then a 0.22 μm syringe filter (Acrodisc, Pall Life Science) to eliminate phytoplankton and bacteria, respectively. 250 mL of the filtered water sample was mixed with 10 mL of sixteen pure bacterial cultures and 10% Zobell Marine Broth (ZMB), and the mixture was incubated at 27°C overnight. Post–incubation, the culture was centrifuged at 8,000 × g for 10 minutes, and the resulting supernatant was filtered using a 0.22 μm syringe filter to remove any residual bacteria. Following the double agar overlay method, plates were incubated at 27°C overnight to observe plaque formation.

### Purification of Phage

For phage adsorption, 200 μL of the filtered sample was mixed with 200 μL of pure bacterial culture in the exponential phase. After the incubation at 27°C for 30 minutes, plaque-forming units (PFU) counts were obtained through the double-agar overlay method. Plaque formation was observed after 12 hours of incubation (11)

To obtain purified phage, a single plaque was picked from the plate and resuspended in 2 mL of SM buffer (2.0 g MgSO_4_·7H_2_O, 5.8 g NaCl,50 mL 1 M Tris-HCl in 1 liter H_2_O, and pH 7.4). The suspension was incubated at 4°C for 2 hours to allow the diffusion. The residual bacteria were removed by filtering the suspension through a 0.22 μm syringe filter. This procedure was repeated 3–4 times to purify the phage. The purified phage stock in SM buffer was maintained at 4°C.

### Multiplicity Of Infection (MOI) of Phage

Briefly, the exponential phase host bacterial culture was mixed with the phage at MOI 0.001, 0.01, 0.1, 1, 10, and 100, and host bacteria without phages served as the control. The mixture was maintained at 27^0^C for 15min to allow phage adsorption and infection. After incubation, the mixture was centrifuged at 8000 g for 5 min. The supernatant was filtered through a 0.22-µm syringe filter to eliminate bacteria. The phage titer was assessed using the double agar overlay method (12). The optimal MOI was determined based on the combination that yielded the highest titer. All the experiments were replicated three times.

### One-step growth curve

A one-step growth curve experiment was conducted to track the replication cycle of the phage by tracing its infection dynamics in the host bacterium (13). The bacterial culture (PA06) in exponential phage (OD_600_ = 0.5–0.7) with optimized MOI (0.1) was mixed with the phage suspension and maintained at 27°C for 10 minutes to improve the phage absorption. Following incubation, the mixture was centrifuged at 6000 x g for 10 minutes, the supernatant was discarded, and the pellet was washed 2 - 3 times with ZMB to ensure the removal of unabsorbed phages. After washing, 50 μL of resuspended cells were added to 50 mL of ZMB and maintained at 27°C for 1 hr. Samples were collected every 10 min for up to 120 min (11). The PFU count was estimated using the double agar overlay method, with three independent repetitions of the experiments.

### Physicochemical stability of phage

To analyse the stability of phage vB_CoeS_P1 in varying ranges of temperature, pH, and salinity, the following experiments were conducted (12). The phage in SM buffer was incubated at varying temperatures ranging from 4°C to 60°C for 1 hr. To examine the influence of different pH levels on phage lysate, the pH of SM buffer was adjusted to a range of 2 -12 using HCl or NaOH, then the buffer was sterilized at 121°C for 20 min. After the sterilization, the phage concentrate was added to the SM buffer and then incubated at 27°C for 1 hr. To assess the phage’s salinity tolerance, SM buffer with varying salt concentrations (0.5%, 3%, 5%, 7%, 9%, 11%, 14%, and 16%) was prepared, and phage concentrate was inoculated. The mix was incubated at 27°C for 1 hr. In all the above-mentioned experiments, the PFU count was measured by using the double agar overlay method.

### Phage Morphology by Transmission Electron Microscopy

The phages in SM buffer were immobilized on 400-mesh carbon-coated grids and centrifuged at 70,000× g for 20 min at 4 °C. After staining, the grids with 2% (w/v) uranyl acetate at room temperature, rinsed twice with 0.02 μm filtered distilled water and dried with filter paper. The slide was observed under a transmission electron microscope operating at 80 kV, with magnification ranging from 20,000 to 60,000×(JEOL 1200Ex, Tokyo, Japan) (14)

### Extraction of phage DNA and Bioinformatics analysis

The phage genome was isolated using Purelink Viral RNA/DNA Mini Kit, following the manufacturer’s protocol (Invitrogen, U.S.A). The isolated genome of the phage was sequenced using the Illumina NovaSeq 6000 platform with 150 bp paired-end sequencing at Humanizing Genomics, Macrogen (Korea).

The raw sequence data were processed and assembled using SPAdes Genome Assembler v3.15.5 (Prjibelski et al., 2020). Before assembly, Trimmomatic v0.39 was used to quality-check and trim the raw paired-end reads (Bolger et al., 2014). Open reading frames (ORF) prediction was accomplished using Pharokka v1.3.2 (Bouras et al., 2023). Predicted ORFs were functionally annotated by using BLASTP against the non-redundant protein database (NCBI) with an E-value threshold of <1e-5, and HHpred, which compared the proteins against four databases: UniProt-SwissProt-viral70_3_Nov_2021, PDB_mmCIF70_16_Aug, SMART_V6.0, and PHROGs_v4 (https://toolkit.tuebingen.mpg.de/tools/hhpred). The terminus of replication and origin in the phage genome were identified through cumulative GC skew analysis. tRNAs encoded by the phage genome were detected using tRNAscan-SE. PhageTerm v1.0.12 was employed to analyze and characterize the phage termini and DNA packaging mechanism (Garneau et al., 2017).

### Phylogenetic tree and comparative genomic analysis

The whole-genome sequence of vB_CoeS_P1 was queried against the NCBI database using tBLASTx with query coverage of 50% and an E-value cut-off of < 1e-5. For finding the evolutionary relationship of vB_CoeS_P1 with other phages, a phylogenetic tree was constructed with conserved genetic elements such as Major Capsid Protein (MCP), and DNA polymerase (DNAP), Terminase large subunit (TerL). BLASTp search against NCBI non-redundant database showing similarity to conserved sequences were selected and phylogenetic relationships were inferred using MEGA 11 software with 1000 bootstrap replicates (15).

To refine the taxonomic position of phage vB_CoeS_P1, a proteomic tree was constructed using the Viptree server with genome-wide similarities computed by BLAST (16). The presence of virulence genes was assessed using the Virulence Factor Database (VFDB) and antibiotic resistance genes by the Antibiotic Resistance Database (ARDB) (Chen et al., 2021; Liu & Pop, 2009). The promoter sequences in the phage genome were identified using the BPROM program, available on the SoftBerry online platform, and putative Rho-dependent transcriptional terminators were predicted using The Arnold program (https://www.softberry.com/berry.phtml?topic=bprom).

Virus Intergenomic Distance Calculator (VIRIDIC) was used to calculate the Average Nucleotide Identity (ANI) and intergenomic similarity between vB_CoeS_P1 and related phages. The results were illustrated as a heatmap (18). To refine the taxonomic position and recognize uncultured homologous phages related to vB_CoeS_P1, the genome was queried against the Integrated Microbial Genomes/Virus (IMG/VR) database with an E-value threshold of < 1e-5, estimated completeness < 50%, and sequence identity > 30% (19). The genomic network analysis of the phage vB_CoeS_P1 and all the screened sequences from IMG/VR, ViPTree, and NCBI databases were examined using vConTACT2 (v0.9.19), in the prokaryotic Viral RefSeq version 201 reference database with ICTV and NCBI taxonomies (20). vConTACT2 tool helps predict the phages’ taxonomic group and classify viral genomes into viral clusters based on the overlapping neighborhood expansion method. The Markov Cluster Algorithm (MCL) identified protein clusters among the genomes. The network map was constructed and visualized using the Gephi (21)

## Result and Discussion

### Phylogenetic Analysis of Bacterial Strain PA 06

The 16S rRNA gene of strain PA 06 revealed 99.65% similarity and 100% query coverage with *Vibrio hepatarius* isolate *UU21* (OX246644.1) when searched against the NCBI BLASTx database. Phylogenetic analysis of the 16S rRNA gene of PA 06 further confirms the close evolutionary relationship with *Vibrio hepatarius* isolate *UU21*, validating the identification of PA 06 as a strain of *Vibrio hepatarius* (Supplementary data S1). The 16S rRNA sequence was submitted to the NCBI repository under the accession number PV177086

### Characterization of vB_CoeS_P1

The phage vB_CoeS_P1 has an optimal MOI of 0.1 (Fig. 1A). The One-step growth curve analysis indicated that the latent period of vB_CoeS_P1 was approximately 20 minutes and the burst size is about 30 PFU/Cell (Fig. 1B). These findings indicate that vB_CoeS_P1 has a comparatively short replication cycle and a moderate burst size, which are typical of many lytic phages.

**FIG 1:**
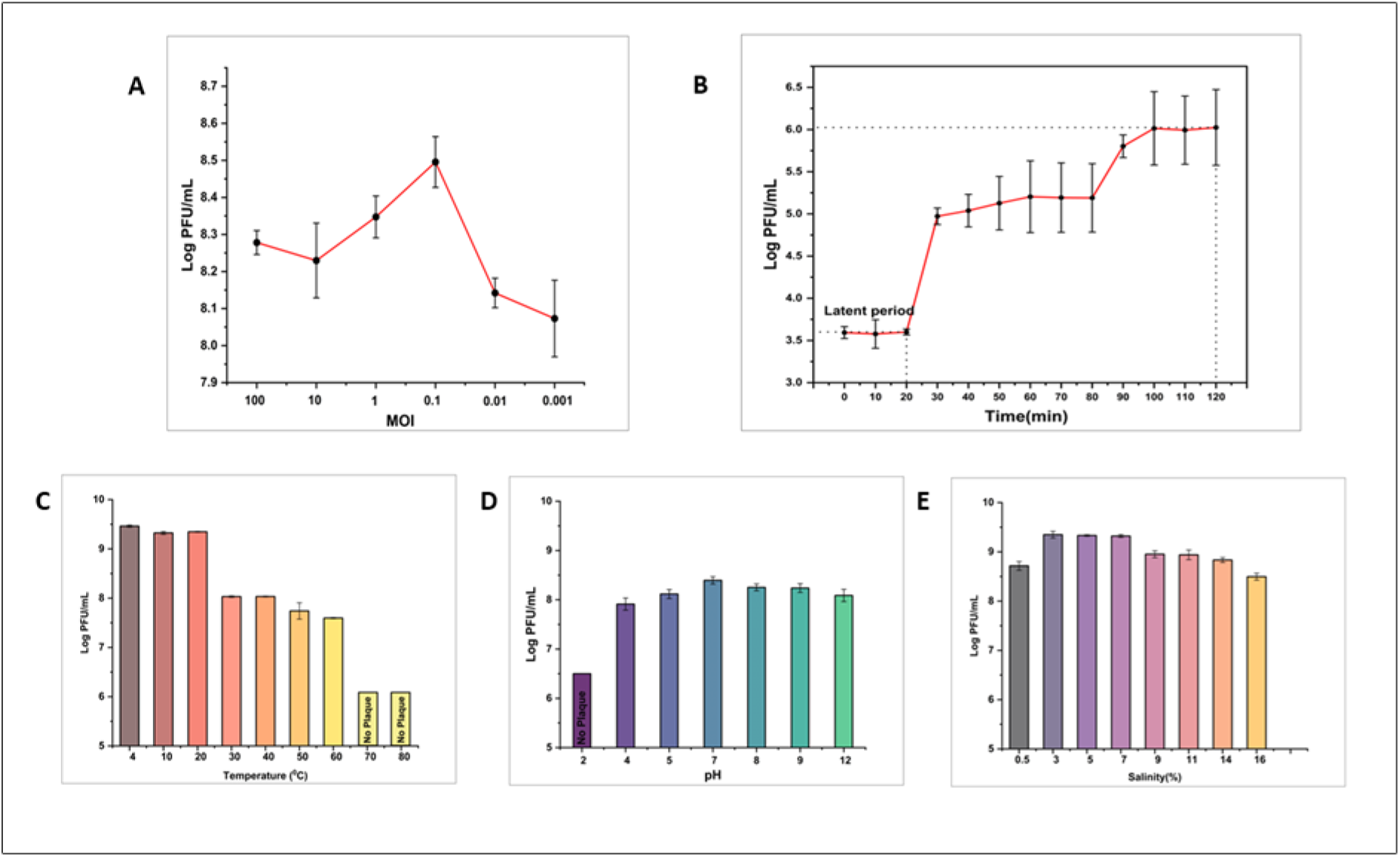
(A) Multiplicity of infection (MOI) graph of vB_CoeS_P1. (B) One-step growth curve of vB_CoeS_P1. The data shown are average values from triplicate experiments, and error bars indicate the standard deviations (SDs). (C) Temperature stability graph of vB_CoeS_P1. (D) pH stability graph of vB_CoeS_P1, (E) Salinity stability graph of vB_CoeS_P1. All the assays were performed in triplicate, and the error bars indicate the standard deviation.

The thermal stability test showed that vB_CoeS_P1 had a wide range of temperature tolerance from 4°C to 60°C. The result showed that the highest survivability rate of 98.14% was in the low-temperature range (4°C,10°C and 20°C) and became inactive at temperatures above 70°C (Fig.1C). This outcome suggests that vB_CoeS_P1 is well adapted to the environment with fluctuating temperatures, which could be advantageous for its survival in natural habitats. The Vibrio phage vB_VpaM_XM1 has a temperature stability profile similar to vB_CoeS_P1; both are stable up to 60 °C and lose activity at 70 °C (22). In contrast, phage vB_VpaP_FE11 was stable between 20 to 50 °C, and there was a drastic decline at 60 °C, and was completely inactivated at 70 °C. Similarly, ValSw3-3, a Siphoviridae bacteriophage infecting *Vibrio alginolyticus*, was stable from 4°C to 50°C but intolerant to higher temperatures (23).). These differences in thermal stability reflect the different ecological niches where the phages and their hosts thrive, such as coastal or estuarine habitats.

pH stability experiments showed that vB_CoeS_P1 is significantly influenced by the varying pH levels. It had an extensive range of pH tolerance from pH 4 to 12. vB_CoeS_P1 shows maximum stability at pH 7 but completely loses its viability at acidic pH 2. However, in alkaline pH 12, the phage survived with a high survivability rate of 98 %, so the phage strongly prefers alkaline environments for survival (Fig. 1D). This adaptation to alkaline pH was also observed in other phages. The phage BP15 infecting *Vibrio parahaemolyticus* demonstrated optimal infectivity rates between pH 6 and 8 and became inactive at pH 2 and 12 (24). In contrast, some phages, like vB_VnaS-L3, showed a narrow range of pH stability (pH 6 to 10), with the highest titer observed at pH 7.5 (25).

Salinity tolerance experiments of vB_CoeS_P1 revealed that it can withstand varying ranges of salt concentrations, with the highest survivability rate (98%) at 3% to 7 % saline concentration. The phage-maintained activity even at high salinity levels of 16% (Fig 1 E). This broad salinity tolerance suggests that vB_CoeS_P1 is highly adapted to estuarine environments, where salinity levels can vary significantly.

### Morphology of vB_CoeS_P1

Phage vB_CoeS_P1 produced small, clear, transparent plaques on the bacterial lawn (Fig. 2A). TEM analysis revealed that the vB_CoeS_P1 is a myovirus-like phage, distinguished by an icosahedral head about 55.5 ± 2 nm in diameter and a long contractile tail extending approximately 97.32 ± 1.3 nm in length (Fig. 2 C and D). The size of the phage was comparable to other phages (Chen et al., 2023; Gu et al., 2024; Lin et al., 2022). Myoviruses are widely abundant and genetically diverse in coastal waters, which shows changes in the community with seasons. A study in the Jiulong River estuary showed that myoviral abundance was higher in near-shore stations compared to offshore, emphasizing that environmental factors influence their abundance and distribution (Liu et al., 2017).

**FIG 2:**
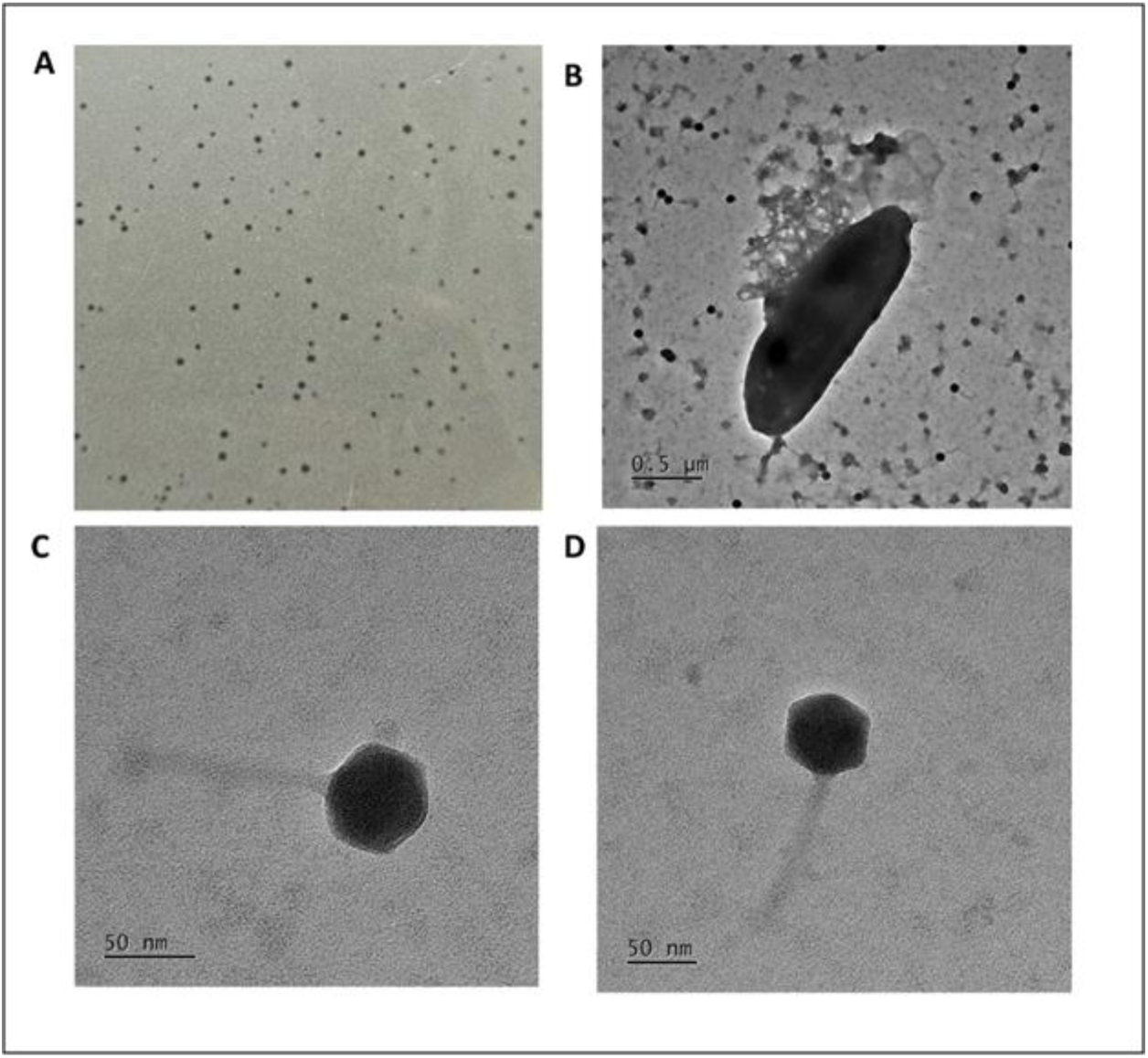
(A) Plaques of vB_CoeS_P1 formed on the bacterial lawn during the double agar overlay method. Transmission electron micrograph of vB_CoeS_P1 (B) Phage released from host cell during the infection (scale bar: 0.5 µm) (Cand D) Single phage with tail (scale bar: 50 µm).

### The Genomic characterization of vB_CoeS_P1

The genome of vB_CoeS_P1 is a double-stranded DNA with 136,927 base pairs, assembled into one single contig with an average of 34.93% GC content (Fig 3). The average genome size of Vibrio phages varies, ranging from 39 kb to 190 kb. Generally, Vibrio phages with myoviral morphotypes have larger genomes (30). The phage vB_CoeS_P1 with 137 kb genome size is comparatively large and closely aligns with the 139,011 bp genome size of phage vB_VpaM_R16F, which infects *Vibrio parahaemolyticus* (Chen et al., 2022). In contrast, the Phage phiTY18 has a much larger genome size of 191,500 bp (Liu et al., 2022). The larger genome size reflects the interplay of ecological, evolutionary, and metabolic factors. It allows the phage to adapt to a wide range of ecological niches, obtain more auxiliary metabolic genes, and improve its evolutionary success (33, 34). Bioinformatic analysis using phageTerm indicated that the vB_CoeS_P1 genome has a circularly permuted structure with redundant ends and utilizes a cos-type packaging mechanism. The vB_CoeS_P1 genome lacks tRNA-encoding genes, highlighting that the phage depends on the host’s translational system for protein synthesis. Phages develop mechanisms like codon usage bias, where the host tRNA codons match with phages, reducing the need for additional tRNAs. For example, the phages JWAlpha and JWDelta, infect multi-resistant *Achromobacter xylosoxidans* and the Delftia lytic phages that infect *Delftia tsuruhatensis* ARB-1, don’t encode any tRNAs. This suggests that the lytic codons in phages are highly adapted to their host and do not require additional tRNAs for translation (35, 36). The Softberry -BPROM promoter prediction software identified 338 potential promoter sites within the vB_CoeS_P1 genome. Analysis of the vB_CoeS_P1 genome through the Comprehensive Antibiotic Resistance Database (CARD) showed no evidence of any antibiotic-resistance genes. Additionally, screening against the Virulence Factor Database did not identify any virulence-related genes in the genome. Notably, several other vibrio phages, such as Artemius phage (infecting *Vibrio alginolyticus*), phi50-12 phage (infecting *Vibrio owensii* GRA50-12), vB_VpP_DE17 phage (infecting *Vibrio parahaemolyticus*), and Siphoviridae phage ValSw3-3 (Infecting *Vibrio alginolyticus*), all of which also lack antibiotic resistance genes and virulence genes (Chen et al., 2020; Droubogiannis et al., 2022; Lin & Tsai, 2022; Yang et al., 2022). The ARNold software predicted 53 transcription terminator sequences within the vB_CoeS_P1 genome. The genome of vB_CoeS_P1 has 265 predicted open reading frames (ORFs), of which 57 were assigned putative functions. The functionally annotated ORFs were designated into five groups: structural proteins, DNA replication, repair and recombination, lysis-related proteins, nucleotide metabolism, and auxiliary metabolic genes-related proteins (supplementary table S1). The whole genome sequence is available in the NCBI database under accession number PV206818.

**FIG 3:**
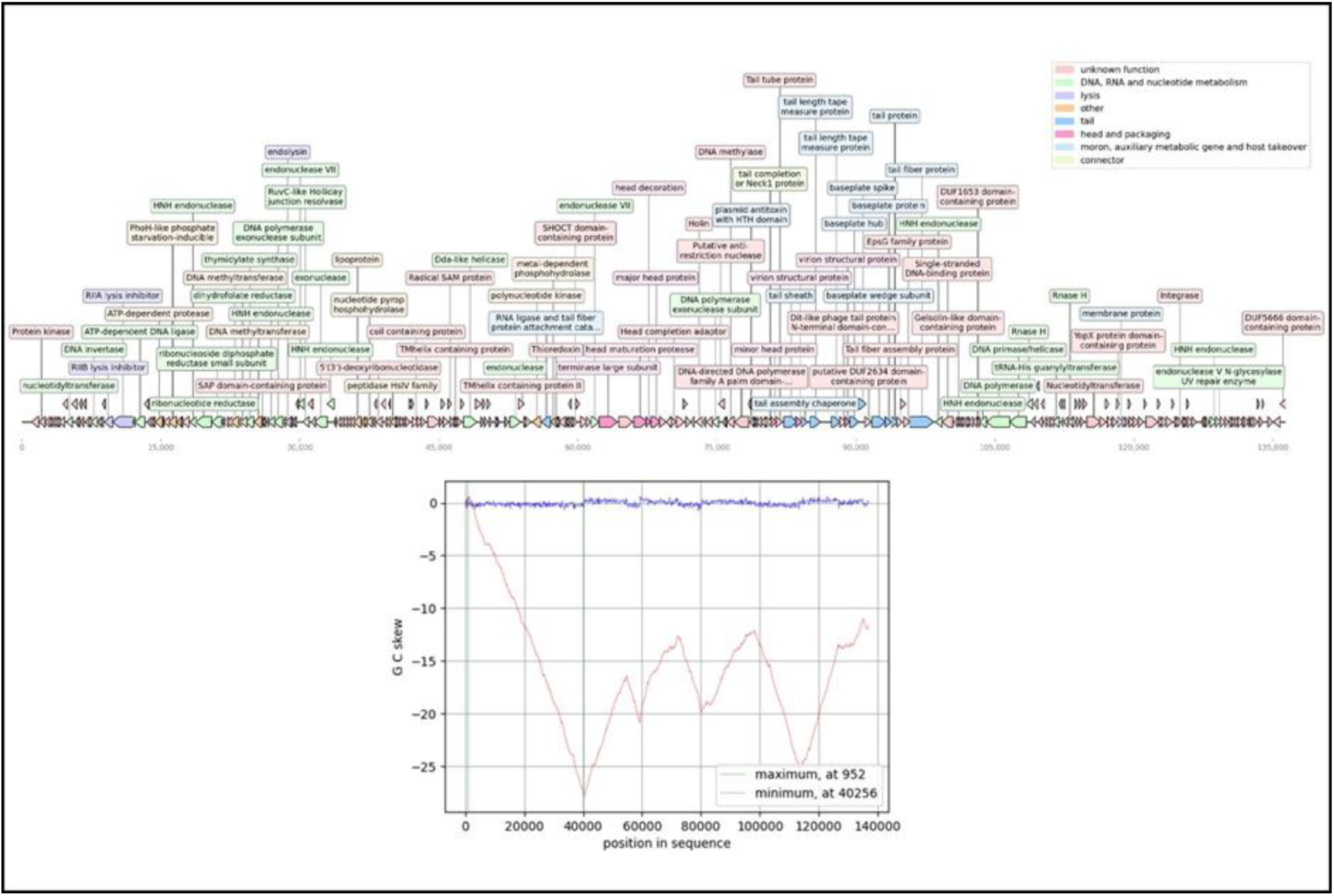
Genomic sequence characterization of vB_CoeS_P1. (A) Genome map of vB_CoeS_P1. Putative functional categories were defined according to annotation and are represented by different colors. (B) Cumulative GC skews of phage vB_CoeS_P1. The global minimum and maximum are displayed in the cumulative graph.

### Structural proteins

Multiple genes were contributed to the vB_CoeS_P1 structural module. The major capsid protein (ORF138) is responsible for constructing the proteinaceous shell that encapsulates and safeguards the phage’s genetic material (40). Phylogenetic analysis showed its highest degree of similarity to the major capsid protein of Vibrio phage vB_VpaM_R16F. ORF138 along with the minor head protein (ORF 163) may also be involved in the viral capsid assembly, serving both structural and scaffolding functions. Head decoration proteins (ORF 137) reinforce the viral capsid, stabilizing its structure and making it more resilient to environmental stresses (41, 42). The head maturation protease (ORF136) is an important enzyme for phage capsid assembly by transforming the precursor proteins into the mature phage head. It is also involved in the successful assembly and functioning of the viral particle (43).

The contractile tail structure of vB_CoeS_P1 is a compound structure, composed of tail sheath protein (ORF 165), tail tube protein (ORF164), baseplate (ORF176), and tail fiber protein (ORF181). During viral infection, the tail fibers and base plate proteins make the initial host recognition and attachment (44). The tail tube is encased within the sheath. After the attachment, the tail sheath contracts in a wave-like manner from the baseplate to the neck (45). This compression drives the tail tube to penetrate the host cell wall and the viral genome is successfully injected (46). Other tail structure-related proteins encoded by vB_CoeS_P1 are tail assembly chaperone (ORF167), tail length tape measure protein (ORF 168), and tail Fiber assembly protein (ORF 178).

The baseplate hub (ORF 172) is a muti-protein structure, from a stable core for the assembly and function of the tail apparatus. It also plays a vital role in initiating the infection process, by host recognition and the initial attachment of the phage to the host surface (47). The baseplate wedge subunit encoded by ORF 175, forms the wedge-shaped segments around the base plate hub (48). ORF173 encoded the baseplate spike protein.

### DNA Replication, Recombination, and Repair

Helicases encoded by ORF 195 help in unwinding nucleic acids during replication. Dda helicase (ORF 105), characterized by its monomeric structure, helps in the unwinding of short DNA duplexes with increased processivity when multiple helicase molecules are on longer substrates (49). The protein encoded by ORF 183, a single-stranded DNA-binding protein (SSB) that protects the single-stranded DNA from degradation during viral replication and repair (Yu & Masker, 2001). The DNA polymerase encoded by ORF 194 exhibits remarkable processivity and fidelity during DNA synthesis. The DNA polymerase exonuclease subunit encoded by ORF 149 possesses 3’ to 5’ exonuclease activity and helps preserve the integrity of the viral genome by enabling its proofreading ability and correcting errors during DNA synthesis. This proofreading function is critical when mutation rates associated with viral replication are high (Zhu, 2014). RNase H (ORF 196) removes RNA primers and resolves RNA/DNA hybrids formed during DNA replication, thereby ensuring the proper processing of viral DNA.

The terminase large subunit encoded by ORF 134 is important for packaging viral DNA into the protein capsid. This subunit facilitates the translocation of viral DNA into the procapsid by ATP hydrolysis. Furthermore, the large terminase subunit with its nuclease activity, cleaves the viral DNA at specific sites and generates free ends for packaging initiation. These critical functions are essential for the assembly of infectious viral particles (52, 53).

### Lysis-related proteins

Holins (ORF146), endolysin (ORF 55), and RIIA/RIIB lysis inhibitor (ORF 24 and ORF 23) proteins work collectively during the host lysis and progeny phage release process. Holins are membrane-bound proteins that create pores in the bacterial cell membrane during lysis (Wang et al., 2000). This triggers the release of endolysins into the periplasmic space and breaks down the bacterial peptidoglycan layer (55). The RIIA and RIIB proteins interfere with holin activity, inhibiting and delaying the lysis process, thereby allowing more time for phage replication and assembly within the host cell (56). Bacteriophage-derived endolysin can be used as alternative therapeutic agent, known as enzybiotics - antibiotic derived from enzymes-against bacterial infections (57, 58). Endolysin CPP-lys from the *Clostridium perfringens* phage has lytic activity against seven different strains of *C. perfringens,* and its applications involve decontamination of fresh produce like lettuce in the food industry (59). Similarly, endolysin from the extremophilic phage phiKo infects the Gram-negative extremophilic bacterium *Thermus thermophilus* HB27, has thermostability and lytic activity against thermophilic bacteria and certain mesophilic bacteria. Additionally, the RAP-29 peptide synthesized from the endolysin has significant antibacterial activity against Gram-positive and Gram-negative bacteria (60). Endolysins, when used in combination with holins and spanins, can enhance their lytic activity and can be used as a potential antibacterial in various fields (61). A green light regulated lytic mechanism in cyanobacterial cultures for biofuel recovery was engineered by introducing T4 bacteriophage-derived holin and endolysin under the control of the cpcG2 promoter. This method enables the energy-efficient recovery of biofuels and related compounds from cells (62).

### AMG – Related proteins

vB_CoeS_P1 encodes Auxiliary Metabolic Genes (AMGs) like pyrophosphohydrolase enzyme (ORF 75), PhoH-like phosphate starvation-inducible (ORF 34), and Endonuclease V N-glycosylase UV repair enzyme (ORF 248). AMG genes carried by the phages can regulate their bacterial host’s metabolism, thereby modulating the nutrient cycles in the marine ecosystem. AMg genes are commonly acquired through Horizontal gene transfer and help to enhance the host metabolic process or improve virus fitness through proteins that are homologous to the host functions (Anantharaman et al., 2018; Enav et al., 2014).

Pyrophosphohydrolase enzyme, a protein product of the mazG gene, hydrolyzes ribo- and deoxyribonucleoside triphosphates into their corresponding monophosphates and pyrophosphate. This activity is crucial in maintaining the nucleotide equilibrium within the host cell. The presence of this gene in the phage genome suggests an evolutionary adaptation to manipulate the host’s nucleotide metabolism, thereby ensuring an adequate supply of nucleotides necessary for viral replication (Gross et al., 2006; Z. Wang et al., 2021).

ORF 34 encoding PhoH-like phosphate starvation-inducible Protein is a critical AMG gene that shows 83.75% similarity and 99% query coverage with Vibrio phage 11895-B1. Approximately 40% of marine phages harbour PhoH genes, making it an effective biomarker. It is a cytoplasmic protein, that binds with ATP and helps control phosphate uptake and metabolism in the phosphate-limited environment (63, 64). Other AMGs associated with phosphorus metabolism include phoB, phoR, pstS, phoA, phoU, phoE, and ugpQ (64)

Homologs of the PhoH gene have been identified among different viral morphotypes with a wide bacterial host range, and have also been found in 40 % of marine phages (63). PhoH is common in cyanophages and Synechococcus phages; P-SSP7, the Podovirus infecting Prochlorococcus phage S-SZBM1, the cyanomyovirus phage S-SRM01 (Rong et al., 2022; Sullivan et al., 2005; Zhang et al., 2021). PhoH has also been documented from other ecological niches. A study in the paddy field in northeast (NE) China showed that more than 400 phages possess PhoH sequences (Wang et al., 2016). Additionally, in northern China, 58 PhoH phage clones were detected from wetland sediments using the primers vPhoHf and vPhoHr (69).

Endonuclease V N-glycosylase, a UV repair enzyme encoded by ORF 248, plays an important role in the phage defense system. It is a bifunctional enzyme with both glycosylase and lyase activities that can initiate the repair of UV-induced pyrimidine dimers. DNA glycosylase initiates the base excision repair pathway by excising pyrimidine dimers by hydrolyzing the glycosylic bond of the 5’ pyrimidine and cleaving the pyrimidine-pyrimidine phosphodiester bond (70). Phages with large genomes often encode endonuclease V, N-glycosylase, and it was well studied in *Escherichia* phage T4 (71). Bacteriophage S-PM2 infects cyanobacterium *Synechococcus* encodes other proteins like UvsX and UvsY for UV-induced DNA damage and repair. A study on two marine *Vibrio parahaemolyticus* hosts and their seven phages shows that host-mediated DNA repair mechanisms, including photoreactivation and excision repair, play an important role in UV-damaged phage DNA (72).

### Nucleotide metabolism-related proteins

The vB_CoeS_P1 genome encodes several important nucleotide metabolisms and tRNA maturation enzymes. tRNA-His guanylyltransferase (ORF 205) plays a crucial role in the maturation of tRNA by adding a guanine nucleotide to the 5’ end of tRNA-His (73). Ribonucleotide reductase comprises two subunits, a larger alpha subunit and a smaller beta subunit, which catalyze the conversion of ribonucleoside diphosphates to their corresponding deoxyribonucleoside diphosphates, essential precursors for DNA synthesis. The smaller beta subunit, encoded by ORF 40, contains a tyrosyl radical-dinuclear iron center believed to start catalysis via electron transfer (74). Dihydrofolate reductase (ORF 46) catalyzes the reduction of dihydrofolate to tetrahydrofolate, a crucial component in amino acid biosynthesis (75). Additionally, the vB_CoeS_P1 virus genome encodes genes for DNA methylase (ORF 153), Nucleotidyltransferase (ORF 10), and 5’-deoxyribonucleotidase (ORF 78).

Some phages with minimal genetic material, like phiX174 with single-stranded DNA (ssDNA) encode only a few proteins and do not contain nucleotide metabolism-related proteins. They rely entirely on the host bacterium for nucleotide synthesis (76). On the other hand, phages with large genomes and complex genetic makeup lack some essential genes. This gap is compensated by host-derived genes or alternative genes from the phage itself. Consider the jumbo phage vB_PaeM_MIJ3 that infects *Pseudomonas aeruginosa* PAO1. It lacks the genes coding for the formation of the phage nucleus, a feature believed to be conserved across jumbo *Pseudomonas* phages (77).

### Phylogenetic and comparative genomic analysis of phage

The NCBI Blastn analysis reveals that vB_CoeS_P1 exhibits very low sequence similarity, with only 1% query coverage and 77.82% identity, to the known phage Vb_VpaM_R16F (OP793884.1). Phylogenetic trees of conserved proteins, including TerL (ORF), DNA polymerase (ORF), and MCP (ORF) were generated using MEGA 11 to reveal the phylogenetic and evolutionary relationship between vB_CoeS_P1 and other related phages, which exhibited the highest similarity in NCBI-BLAST analysis. The results revealed that, for MCP and DNA polymerase, vB_CoeS_P1 clustered with vB_VpaM_R16F, forming a distinct clade. In contrast, for TerL, vB_CoeS_P1 did not cluster with any other phages and instead formed a unique clade, highlighting its divergence from other known phages (Fig 4).

**FIG 4:**
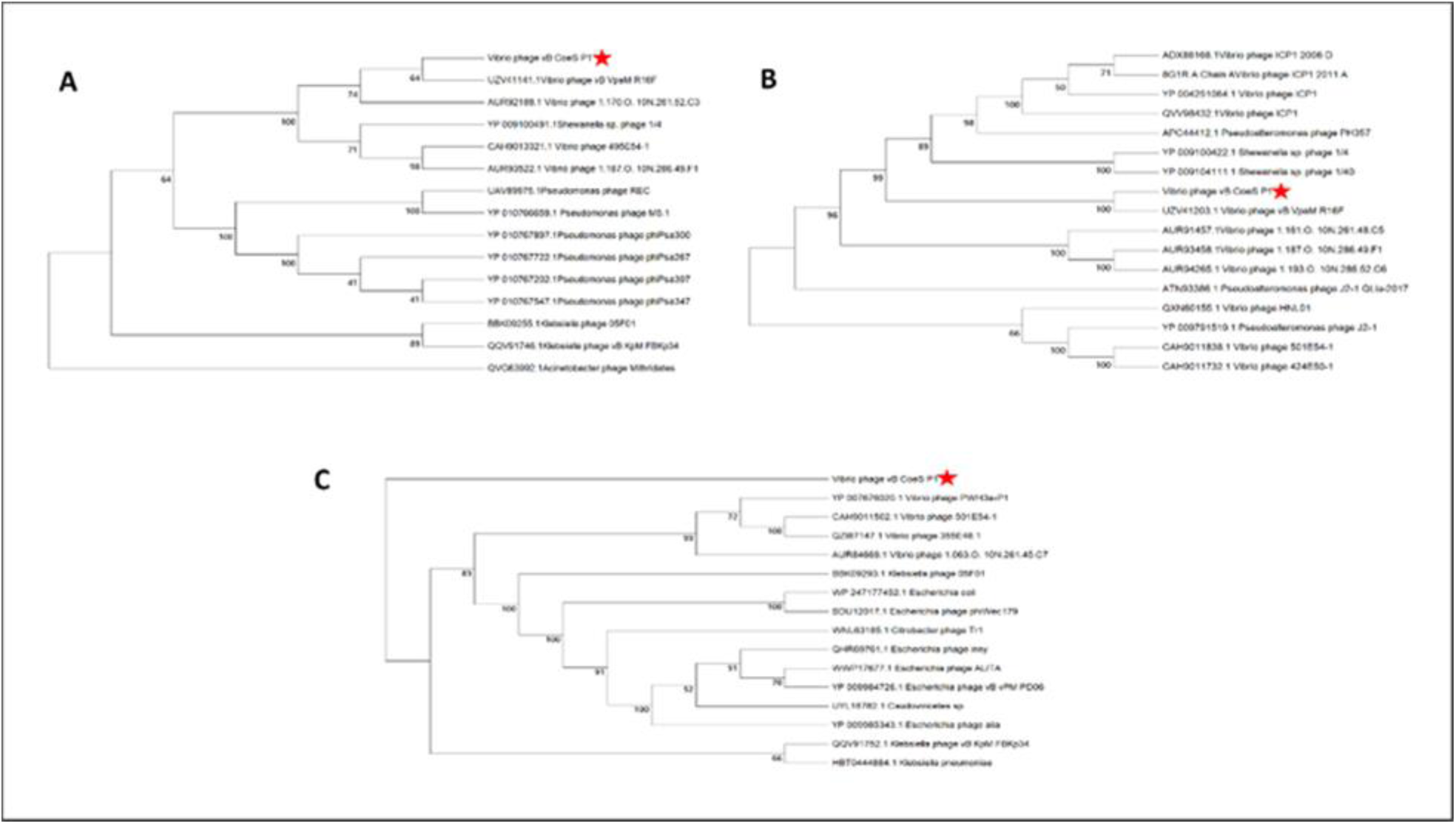
Phylogenetic trees based on the terminase large subunit (A), major capsid proteins (B), and the DNA polymerase (C) of vB_CoeS_P1 and homologous proteins from other similar phages. Amino acid sequences were aligned using MEGA 11 software, and the phylogenetic trees were generated using the neighbor-joining method with 1,000 bootstrap replications.

Using Viptree, a circular proteomic tree was constructed to analyze the phylogenetic relationships between vB_CoeS_P1 and other reference phages in the database. A circular proteomic tree was constructed using 4,473 selected phages (Fig. 5A). Additionally, 18 phages exhibiting the highest similarity to Vb_VpaM_R16F were shortlisted to develop a rectangular proteomic tree (Fig 5 B). The phylogenomic analysis based on proteomic data indicates that vB_CoeS_P1 forms a distinct evolutionary lineage, divergent from other known phages. To corroborate this finding, a comparative genomic analysis was performed on the most closely related phages using Viptree DiGAlign, with six (NC0208631, NC254701, NC0254361, NC0290571, NC0478391, NC0208431) complete phage genome sequences selected based on tBLASTx comparisons. Comparative genomic analysis results revealed that vB_CoeS_P1 has a low degree of genomic similarity to the other examined phages (Fig 6).

**FIG 5:**
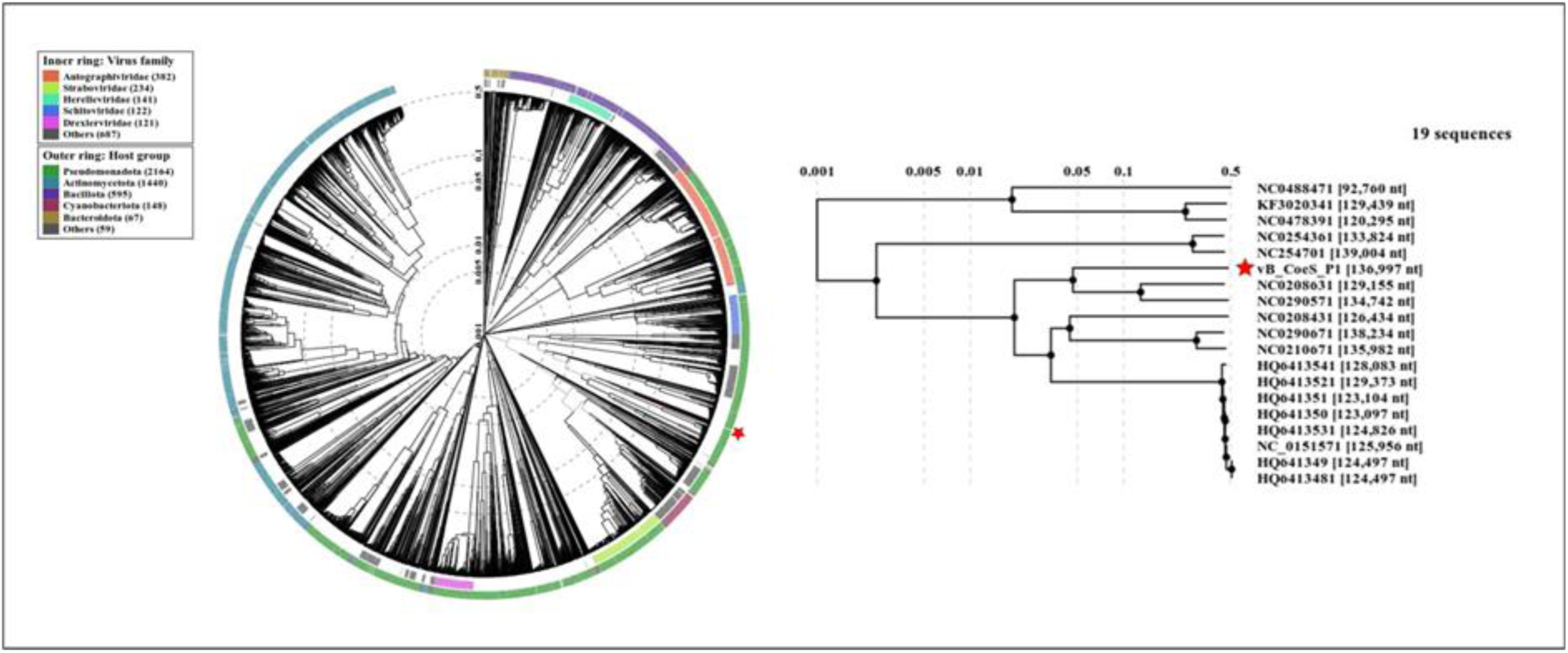
Proteomic tree of vB_CoeS_P1 phage. (A) ViPTree circular proteomic tree of prokaryotic dsDNA viruses with vB_CoeS_P1. vB_CoeS_P1 phage sequence is highlighted with red stars. (B) The rectangular presentation of the proteomic tree shows the closest related phages to vB_CoeS_P1.

**FIG 6:**
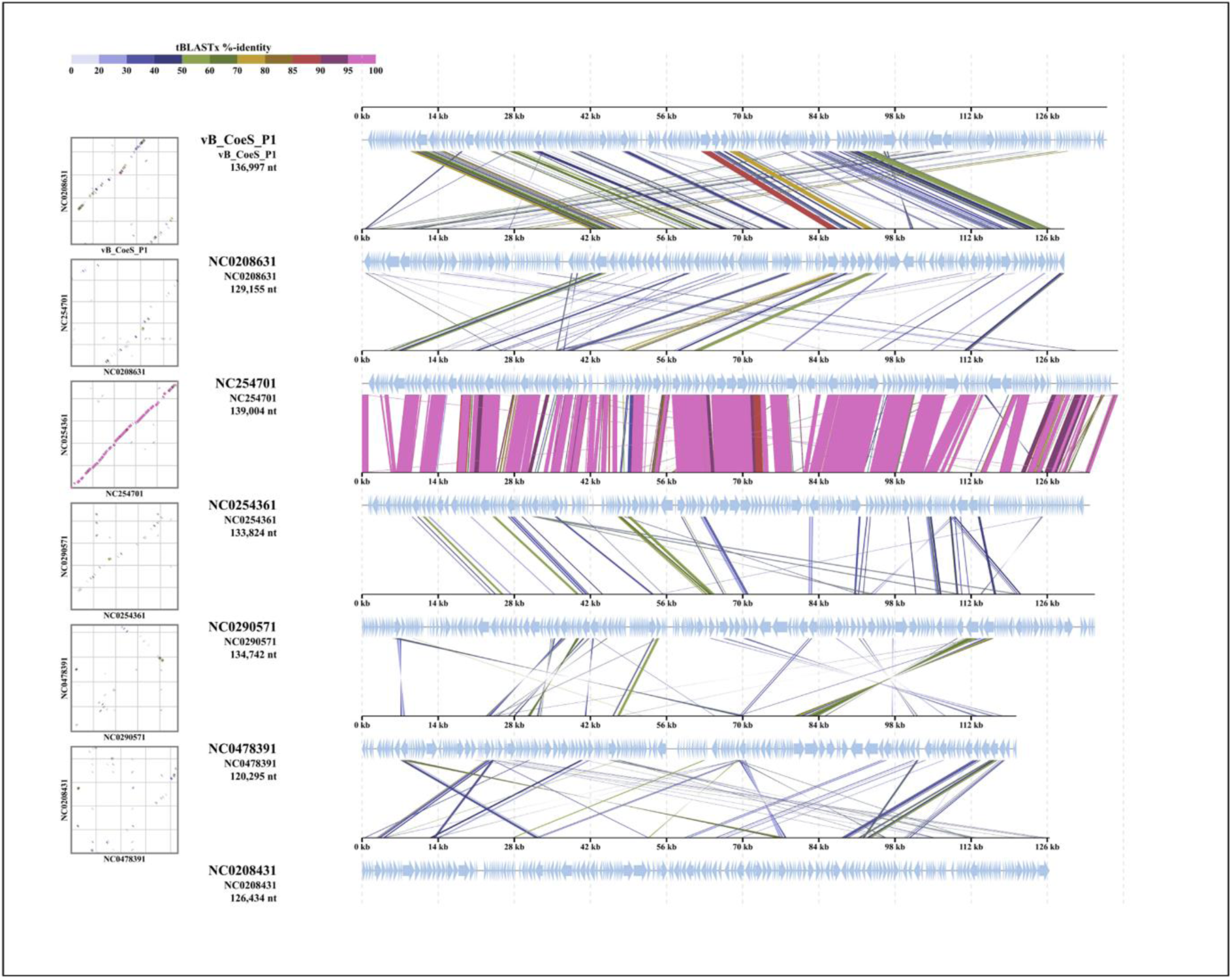
Genomic comparisons between vB_CoeS_P1 and its related phages. The shading below each genome indicates sequence similarities between the genomes, with different colors representing the levels of similarity.

Based on the comparative genomic analysis results, vB_CoeS_P1 appears to have the potential to represent a new family. However, it is challenging to establish this finding conclusively with a single phage isolate. To better understand the evolutionary relationships between vB_CoeS_P1 and other uncultured phages, an all-versus-all BLAST search was conducted using the Integrated Microbial Genomes and Viruses (IMG/VR) database. The analysis revealed that 12 metagenomically assembled, uncultured viral genomes displayed similarity to vB_CoeS_P1 (supplementary table S2).

A Nucleotide – based intergenomic similarity analysis was conducted using the VIRIDIC tool to evaluate the relationships between vB_CoeS_P1 and other phages showing similarity, including 12 from the IMG/VR reference database, 14 from NCBI, and 19 from Viptree. The results indicated that the nucleotide-based intergenomic similarity between vB_CoeS_P1 and other phages ranged from 0.2 to 9.2%. The ICTV subcommittee has established Average Nucleotide Identity (ANI) thresholds for delineating viruses into species (95%) and genera (∼70%) (18) (Fig 7). Given the very low ANI observed, vB_CoeS_P1 is notably distinct from other isolated phages and is proposed as a representative of a new family within the *Caudoviricetes* class.

**FIG 7:**
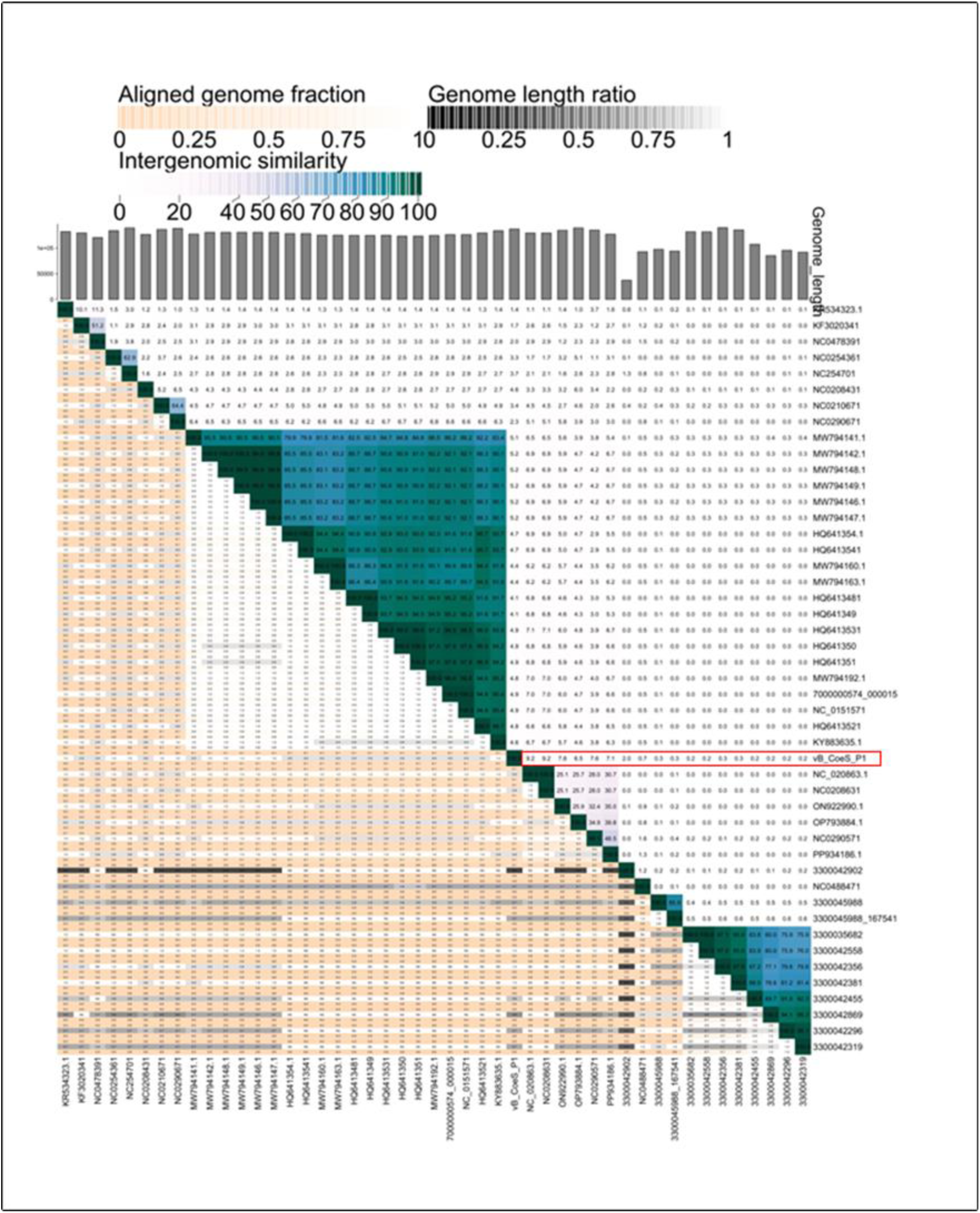
Heat map of intergenomic similarities (right half) between vB_CoeS_P1 and its distantly similar phages. Values represent Average nucleotide identity (ANI) generated by VIRIDIC.

The network analysis using vContact2.0 reveals that vB_CoeS_P1 does not belong to any established viral clusters but rather appears to represent a potential outlier group with very low similarity to the reference databases. Furthermore, the vContact2.0 taxonomic prediction indicates that vB_CoeS_P1 and most of the phages displaying genomic similarity with it from various databases are classified within the unassigned group, suggesting that the taxonomic family of these phages remains undetermined (Figure 8). Therefore, we can propose that vB_CoeS_P1 represents a new family within the *Caudoviricetes class,* named here as *Estoviridae*

**FIG 8:**
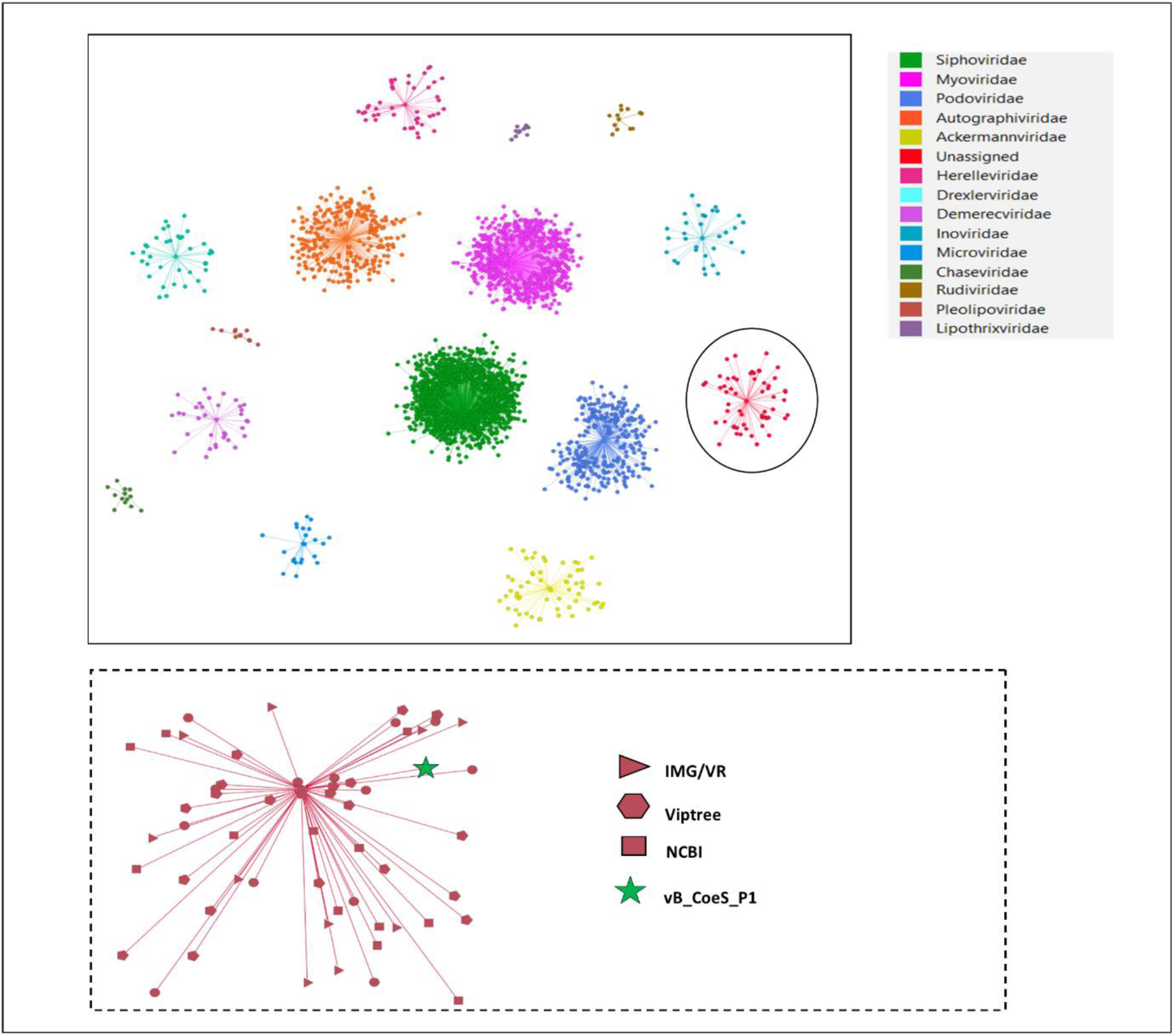
A network analysis of vB_CoeS_P1 and other distantly related phage from Vipree, NCBI, and IMG/V by vConTACT2. The nodes represent the viral genomic sequences, and the different colors of the nodes represent different viral families.

## Conclusion

In marine ecosystems, viruses hold a vital role in shaping microbial communities and driving biogeochemical cycles. In this study, a novel phage vB_CoeS_P1 was isolated from the coastal water of Cochin Estuary, representing the first known phage for *Vibrio hepatarius*. vB_CoeS_P1 demonstrates stability across a broad thermal range, salinities, and pH levels, along with a lytic life cycle and a burst size of 30 PFU/mL. This result indicates that vB_CoeS_P1 has adaptive features that help to thrive in dynamic coastal environments. The presence of AMGs further supports the survival of both host and phage under varying environmental conditions. The whole genome shows that vB_CoeS_P1 has a genome of 136,927 base pairs, encoding 265 predicted ORFs, of which 57 were assigned putative functions. Phylogenetic analysis revealed the unique evolutionary lineage of vB_CoeS_P1, positioning it as a representative of a new viral family *Estovirdae* within the *Caudoviricetes* class. This study provides a holistic genomic and phylogenetic characterization of the first known phage of *Vibrio hepatarius,* offering valuable information about host-phage interactions in the marine environment.

## Data Availability

The 16S rRNA sequence of PA05 and the whole genome sequence of vB_CoeS_P1 have been deposited in the NCBI database with accession numbers PV177086 and PV206818.

## Acknowledgment

The authors acknowledge the Head of the Department, Department of Biotechnology, Cochin University of Science and Technology, for providing the necessary research facilities. The authors also express their gratitude to the Vice Chancellor of CUSAT for institutional support. This study was carried out as part of a project funded by the Science and Engineering Research Board (SERB). The authors are grateful to Sritha K. S. for her valuable suggestions in data analysis. Additionally, the authors extend their gratitude to the Council of Scientific and Industrial Research (CSIR) and the University Grants Commission (UGC), New Delhi, for providing research fellowship grant.

## Conflicts of Interest

The authors declare no conflicts of interest.

## Author Contributions

Shruthi P K - Conceptualization, Writing -original Draft, Data curation, Investigation, Methodology, Software, and Visualization

Parvathi Ammini – Conceptualization, writing, Review and Editing, Funding acquisition, Supervision, Data curation, Methodology

Pradeep Ram Angia Sriram – Methodology and Visualization

## Funding Agency

This study formed a part of the project: DST-SERB-Marine microbiome to ascertain the role of microbes in biogeochemical cycling in the Eastern Arabian Sea (SRG/2022/001545)

